# PhenoComb: A discovery tool to assess complex phenotypes in high-dimension, single-cell datasets

**DOI:** 10.1101/2022.04.06.487335

**Authors:** Paulo E. P. Burke, Ann Strange, Emily Monk, Brian Thompson, Carol Amato, David M. Woods

**Affiliations:** Division of Medical Oncology, Department of Medicine, University of Colorado School of Medicine, Aurora, CO 80045, USA

## Abstract

**Motivation:** High-dimension cytometry assays can simultaneously measure dozens of markers, enabling the investigation of complex phenotypes. However, as manual gating relies on previous biological knowledge, few marker combinations are often assessed. This results in complex phenotypes with potential for biological relevance being overlooked. Here we present PhenoComb, an R package that allows agnostic exploration of phenotypes by assessing all combinations of markers.

**Design:** PhenoComb uses signal intensity thresholds to assign markers to discrete states (e.g. negative, low, high) and then counts the number of cells per sample from all possible marker combinations in a memory-safe manner. Time and disk space are the only constraints on the number of markers evaluated. PhenoComb also provides several approaches to perform statistical comparisons, evaluate the relevance of phenotypes, and assess the independence of identified phenotypes. PhenoComb allows users to guide analysis by adjusting several function arguments such as identifying parent populations of interest, filtering of low-frequency populations, and defining a maximum complexity of phenotypes to evaluate. We have designed PhenoComb to be compatible with local computer or server-based use.

**Results:** In testing of PhenoComb’s performance on synthetic datasets, computation on 16 markers was completed in the scale of minutes and up to 26 markers in hours. We applied PhenoComb to two publicly available datasets: an HIV flow cytometry dataset (12 markers and 421 samples) and the COVIDome CyTOF dataset (40 markers and 99 samples). In the HIV dataset, PhenoComb identified immune phenotypes associated with HIV seroconversion, including those highlighted in the original publication. In the COVID dataset, we identified several immune phenotypes with altered frequencies in infected individuals relative to healthy individuals. Collectively, PhenoComb represents a powerful discovery tool for agnostically assessing high-dimension, single-cell data.

**Availability:** The PhenoComb R package can be downloaded from https://github.com/SciOmicsLab/PhenoComb

## 1 Introduction

Flow and mass cytometry assays are able to measure dozens of markers in hundreds of thousands of cells per sample at single cell resolution[1], enabling the characterization of complex phenotypes and powering new insights. However, traditional, hypothesis-driven, manual gating analysis of flow data relies on prior knowledge of the researcher, enabling only incremental advances. In contrast, agnostic approaches to the analysis of flow data are powerful discovery tools that more fully utilize the high-dimension data and are not limited to *a priori* hypotheses. However, many agnostic approaches, such as dimension reduction *(e.g.* UMAP[2]) and clustering tools *(e.g.* Phenograph[3]), fail to capture discrete phenotypes, limiting interpretability, the ability to do statistical analysis, and downstream isolation of cells *(e.g.* flow sorting) for future experiments.

An alternative agnostic approach is to exhaustively identify discrete phenotypes in the flow data by assigning discrete states for each of the measured markers and counting the cells that have the combination of markers states of a given phenotype. For example, if two markers (A and B) are measured and discretized into positive (+) and negative (-) states, an exhaustive combination of markers and states would result in the following phenotypes: A+B+, A+B-, A-B+, A-B-. Additionally, the partial combinations of markers could also be considered: A+, A-, B+, B-. By visiting all possible marker/state combinations, the cell counts could be then used to assess which phenotypes are relevant given a comparison of interest.

Similar approaches were shown to be effective, unveiling novel phenotypes in the context of cancer therapy[4] and infectious diseases[5]. However, exhaustive exploration in the phenotypic landscape can rapidly get computationally expensive to perform as the number of markers increases. This is because the number of possible phenotypes increases exponentially, such as (*n* + 1)^*m*^ – 1, with the number of markers (*m*) and states (n). Previous approaches could only tackle a limited number of markers, had long run-times, and are no longer supported/available. Thus, new and more powerful bioinformatic tools are needed to perform this kind of analysis.

Here we present PhenoComb, an efficient tool for the exploration of complex phenotypic landscapes. It is available as an R package and implements methods to 1) convert continuous scale data for marker expression into discrete states, 2) count cells for all possible phenotypes present in the input dataset, 3) to perform a variety of statistical comparisons to evaluate the relevance of those phenotypes based on cell frequency of the samples, and 4) to identify relevant phenotypes that are independent from each other. PhenoComb takes advantage of parallel computing and is implemented in an memory-safe manner, overcoming limitations on the number of markers from previous approaches. It can process datasets with virtually any number of markers, only being limited by computing time and disk space for the outputs.

## 2 Methods

### 2.1 Package Implementation and Features

PhenoComb is available as a R package with some functions implemented in C++ for efficiency, but only available through R’s interface. There are two coding workflows available that implement the analysis illustrated in Figure 1: a local workflow, designed for ease of use, holding all objects in memory, being easily accessible for the user; and a server workflow, designed to handle large datasets which can’t be held in memory. In both implementations, PhenoComb takes advantage of parallel computing to increase performance where steps of the analysis are mutually independent. Tutorials of how to install and use PhenoComb are available at the GitHub repository page: https://github.com/SciomicsLab/PhenoComb.

**Figure 1:**
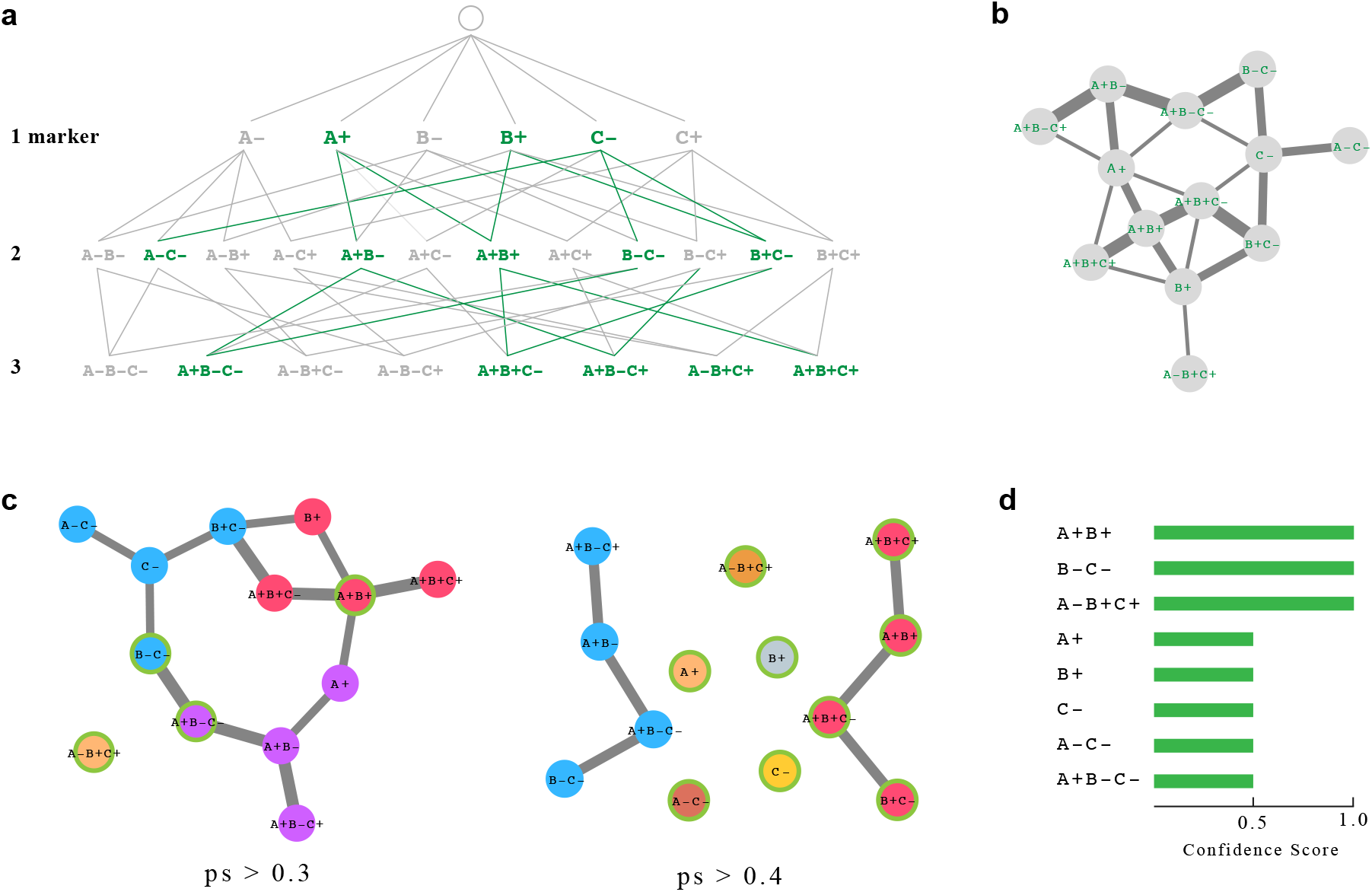
PhenoComb analysis outline. **(a)** Given a set of markers, all possible marker state combinations with lengths from one to the number of markers are generated, each one being considered a unique phenotype. Cells matching each phenotype are counted for each sample. A statistical test is performed comparing phenotype frequencies between two groups of samples and the ones meeting a defined significance threshold are selected. In the illustrated hypothetical, selected phenotypes are shown with a darker color. **(b)** A phenotypic similarity score (ps) is computed between all selected phenotypes and represented as a network. The thickness of the edges indicates their similarity. **(c)** The network has its edges filtered by several ps thresholds and a clustering algorithm is applied for each one. For each cluster identified, the phenotype with the highest log2fold-change is selected as the representative one. Clusters are represented by different colors and representative phenotypes are highlighted with a green border. **(d)** A confidence score for each phenotype is then computed as the frequency of its selection as a representative throughout the networks.

The package also provides several ways to filter data and tune the analysis. For example, cells can be filtered out if intensity values are over a certain threshold, called Out of Bound (OOB), to reduce noise. When counting cells for all phenotypes, the user can filter phenotypes by the minimum number of cells on a fraction of samples, by parent phenotype, and also set the maximum phenotype length desired. For the statistical tests, the output can be filtered by parent population and p-value. PhenoComb also provides some convenience tools to deal with big outputs, such as finding specific phenotypes in files.

### 2.2 Combinatorial Phenotypes Computation

Given a dataset with *m* markers’ signal intensity measured, each continuous measurement can be discretized into a finite number *n* of states through *n* — 1 provided thresholds obtained by manual gating. These states can be called, for example, negative (-) and positive (+) if one considers only two states, or negative (-), low (+), and high (++) if three states are considered. The number of discrete states can be any number equal or greater than two, and there can be a different number of states for each marker.

Considering that all m markers can be discretized into n states, there are *n^m^* possible combinations of marker states. Each different combination will henceforth be referred to as phenotypes. Additionally, we can consider phenotypes that are neutral for one or more markers, increasing the total number of possible phenotypes to (n + 1)^*m*^ – 1.

To illustrate the combinations between markers and states, Figure 1a shows all possible combinations between three markers (A, B, and C) each one having two possible states (- and +). Each row indicates the number of markers for the phenotype, henceforth called length, as a consequence of considering phenotypes neutral to certain markers.

PhenoComb provides a method to count the number of cells in the dataset that have a given phenotype for all possible phenotypes across all samples. This is done efficiently by first counting cells for all full-length phenotypes present in the data, and then generating all other phenotypes with neutral states by summing up the cells already counted in the full-length ones. Since the computation of each neutral-state combination is independent from each other, this step can take advantage of parallel computation to increase performance.

### 2.3 Evaluation of Generated Phenotypes

The statistical evaluation of a phenotype based on its cell counts across samples can be done in several ways. PhenoComb provides three types of statistical comparisons: two group comparison using the Man-Whitney U test, correlation with a given variable using Kendall’s rank correlation, and a time-to-event (survival) analysis test using Cox Proportional Hazards Model. Since total number of cells can vary across samples, all statistical tests are performed on cell frequencies, being the cell count of the phenotype divided by the total number of cells the sample contains. If phenotypes are filtered for a given parent phenotype, the frequencies will be calculated based on the parent phenotype’s cell count instead of total number of cells. The phenotypes can then be filtered by respective p-values as represented in bold font on Figure 1a.

While an exhaustive evaluation of marker combinations can create a large number of phenotypes, these phenotypes are not independent of one another. For example, in a two marker dataset, the phenotype A+ is related to the phenotypes A-, A+B-, and A+B+. Given the interrelatedness of phenotypes, when performing statistical comparisons, the type I error rate (i.e. false positives) is not a product of the number of comparisons. Therefore, traditional methods for correcting for multiple comparisons are not appropriate. However, an inflated type I error rate (i.e. false positives) will still occur. PhenoComb does not adjust for multiple comparisons, and should be used with that caveat.

### 2.4 Identification of Independent Phenotypes

As mentioned in the previous section, phenotype cell counts can be dependent on each other. In large datasets, it can be hard to distinguish the phenotypes that are the actual drivers of statistical/biological relevance. PhenoComb provides a network-based algorithm aimed at finding the driver phenotypes.

Given a number *P* of phenotypes, a phenotypic similarity (*ps*) score matrix *S_P×P_* is calculated based on the number of markers each pair of phenotypes share, such as:

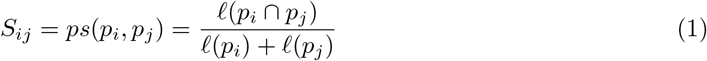

where *ℓ* is the length of a given phenotype. The similarity matrix *S* can be understood as a symmetric adjacency matrix, thus representing a undirected network where each node is a phenotype, and the links between them are weighted by the *ps* score given in Equation 1. An example of such network is depicted in Figure 1b.

Next, *n* sub-networks are generated by pruning the original network from its edges by applying a set increasing thresholds *t*_1_,*t*_2_,…,*t_n_* with

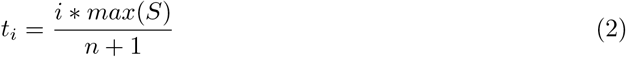

The pruning is performed by applying the following function to the matrix *S*:

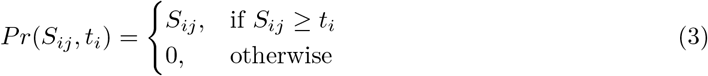

For each sub-network, we apply the Louvain clustering algorithm[6] which automatically finds the optimal number of clusters and labels the nodes accordingly, as illustrated in Figure 1c. A representative phenotype is then selected from each cluster by ranking them according to the absolute value from the statistic provided by the chosen comparison. For group comparison, the ranking value used is the |effect size| * |*log*_2_(fold change)ļ. We use this formula to avoid artifacts such as only one sample with an unexpected high frequency driving the fold change. For the correlation test, the correlation metric is used. The Cox Proportional Hazards coefficient is used if this method was chosen. All values used for ranking are normalized for the range. Finally, the representative phenotypes are ranked by a confidence score based on the frequency that the given phenotype shows up as representative in the *n* sub-networks, as depicted in Figure 1d.

### 2.5 Datasets

#### 2.5.1 Synthetic Datasets

To better evaluate PhenoComb’s performance, we generated a set of synthetic datasets simulating several combinations of number of markers, maximum phenotype length, number of cells, and number of samples. Since they were only used to test the performance of the combinatorial phenotypes computation, all full length phenotypes for all samples have a similar number of cells. The code to generate such datasets is available on the aforementioned GitHub repository. All computations were performed on a 28-core, 56-thread, Intel Xeon W-3275M 2.50GHz with 1TB of memory. However, the memory requirements for PhenoComb are approximately four times the size of the fcs input file.

#### 2.5.2 The HIV Dataset

This dataset, previously described in Weintrob *et al.* [7], was obtained from FlowRespository.org, flow repository ID FR-FCM-ZZZK. The dataset consisted of 466 PBMC samples obtained from HIV infected military personnel that were evaluated by flow cytometry for 12 markers (CD3, CD4, CD8, CD45RO, CD27, CD28, CD57, CCR5, CCR7, CD127, KI-67 and CD14) and a viability dye. Samples contained a mean of 393,315 events. Individual sample fcs files were gated to exclude debris (FSC-A vs. SSC-A). Samples with less than 5000 cells were not considered, leaving a total of 421 samples to be analyzed. Dead cells and CD14+ cells were removed through a dump channel gate. Samples were then concatenated and each marker gated as in Supplemental Figure 1A. These analyses were performed in FlowJo 10.7 software. The fluorescence intensity coordinates of gates (i.e. threshold differentiating positive and negative populations for each marker) were recorded in a csv file for reference in PhenoComb function arguments.

#### 2.5.3 The COVIDome Dataset

This dataset, previously described in Sullivan *et al.*[8], was also obtained from FlowRespository.org, flow repository ID FR-FCM-Z367. The dataset consisted of 99 PBMC samples obtained from COVID infected individuals (n=69) and COVID negative individuals (n=30) which were evaluate for 40 markers by CyTOF mass cytometry. Samples contained a mean of 390,059 events. The publicly available fcs files were previously gated to remove anomalies and dead cells. Sample fcs files were concatenated, each marker gated as in Supplemental Figure 1B, and the intensity coordinates of gates were recorded in a csv file.

## 3 Results

### 3.1 Performance Benchmark

To evaluate PhenoComb’s performance, we simulated a set of 48 synthetic datasets considering different numbers of markers, cells, samples, maximum phenotype length, and number of threads used to compute. These datasets represent the worst-case scenario for each configuration of parameters since all possible combinations of full-length phenotypes will be present in them. That is unlikely to happen in real datasets because of biological restrictions and limited number of sampled cells. To illustrate the magnitude of the analysis, Figure 2a shows the number of expected computed phenotypes in the synthetic datasets for a given number of markers. It shows the exponential nature of the combinatorial problem. Also, the number of phenotypes in HIV and COVIDome datasets (discussed later) is shown in purple, confirming that the number of phenotypes present in real datasets is lower than on the theoretical ones, despite some markers in the real datasets having more than two states.

**Figure 2:**
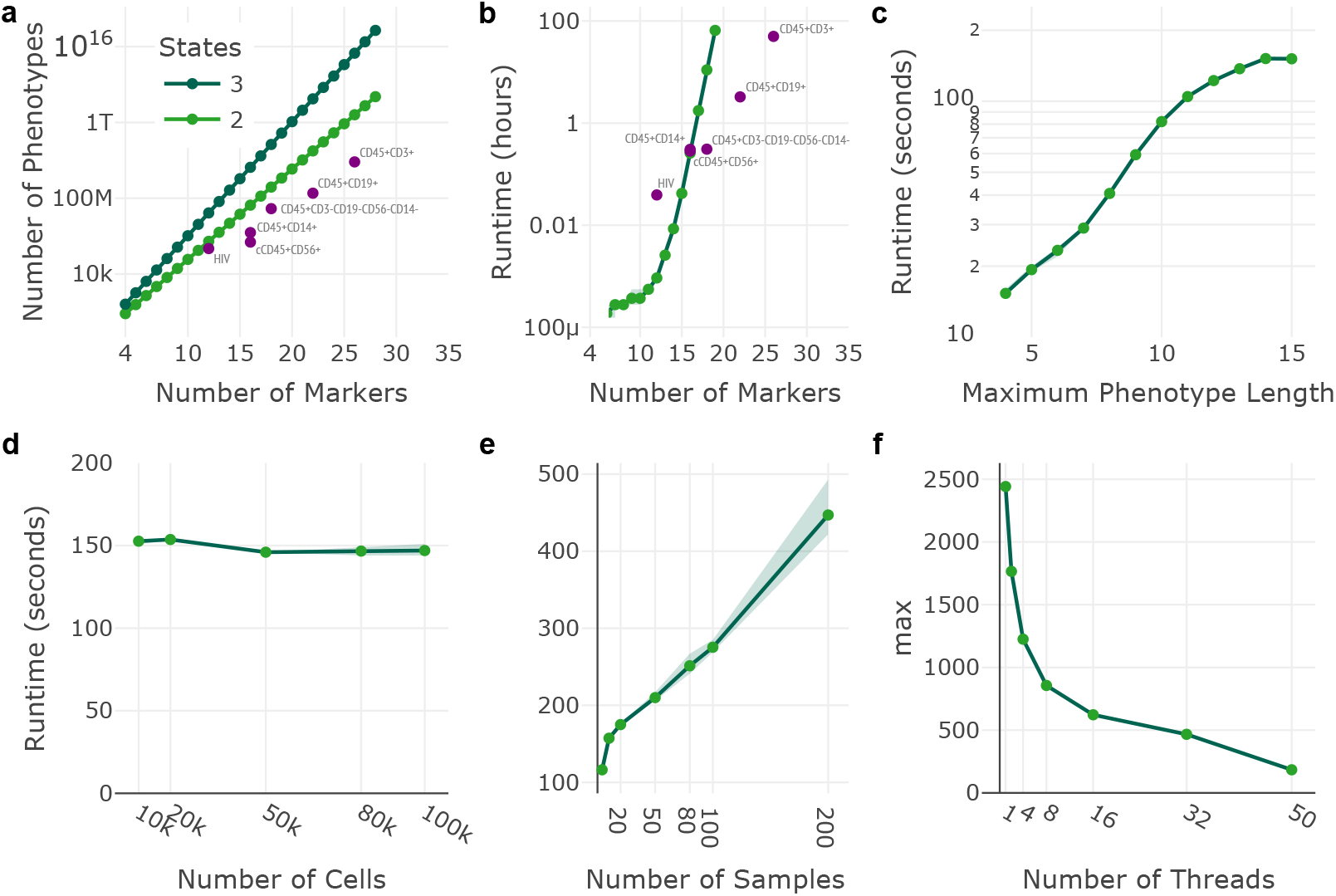
Performance Benchmark. Computing times for the combinatorial phenotype generation step are shown in green for synthetic datasets and in purple for the HIV and COVIDome datasets. a) The number of phenotypes generated by increasing numbers of markers. b) The amount of time (hours) to analyze various marker sizes. c) The amount of time (seconds) to analyze various maximum phenotype lengths. d) The amount of time (seconds) to analyze various number of cells. e) The amount of time (seconds) to analyze various numbers of samples. f) Computing time (seconds) reduction by increasing the number of threads.

For each of the experiments shown in Figures 2b-f, we varied one parameter and fixed the remaining parameters as follow: 15 markers, 40,000 cells per sample, 10 samples, 50 threads, and the maximum phenotype length as the number of markers unless otherwise stated. All datasets were assessed with 2 states for each marker. All analysis were performed using the server workflow since the local workflow only allows the processing of small datasets, which are usually computed in a matter of a few minutes. The experiments shown in Figure 2 account for the combinatorial phenotype counting step, which is the most impacted by the number of markers and states.

The parameter that impacts PhenoComb’s performance the most is the number of markers present in the dataset. This was expected since the number of possible phenotypes grows exponentially with the number of markers. Figure 2b shows that the computational complexity growth is super-exponential. PhenoComb allows a decrease in the maximum length of phenotypes to mitigate this problem when very large phenotypes are not of interest. This feature can also decrease exponentially the computation time as shown in Figure 2c. Nevertheless, computation times for real datasets are almost always lower then the synthetic ones because of the lower number of present phenotypes, even though the number of samples and the cell counts per sample were higher in the real datasets used vs. the synthetic datasets. The only case where the real dataset is higher than the synthetic is for the HIV data because of its high number of samples (n=421).

The total number of cells in the dataset had little impact on the cell counting process once they are all transformed to single numbers when calculating the full-length phenotypes, a relatively computationally inexpensive process (Fig. 2d). The number of samples present in the dataset increase linearly the time to compute the phenotypes as shown in Figure 2e. Lastly, PhenoComb can take advantage of parallel computing, reducing almost exponentially the computing time as the number of threads used is increased (Fig. 2f).

### 3.2 Validation of Phenotypes Identified in Previous Approaches

To test PhenoComb’s ability to identify biologically meaningful phenotypes, we used a previously published, publicly available flow cytometry dataset evaluating T-cell immunophenotypes associated with HIV seroconversion[5]. PhenoComb was utilized to discretize markers and assess all marker combinations. Marker combinations for CD3+ population were then assessed for associations with survival time following seroconversion. 9754 phenotypes with an unadjusted p-value of less than 5.0E — 7 were identified. We specifically focused on phenotypes identified in the original publication (Table 1). Of the three phenotypes highlighted in the original publication as being significantly correlated with time to seroconversion, PhenoComb identified two as also statistically significant.

**Table 1:**
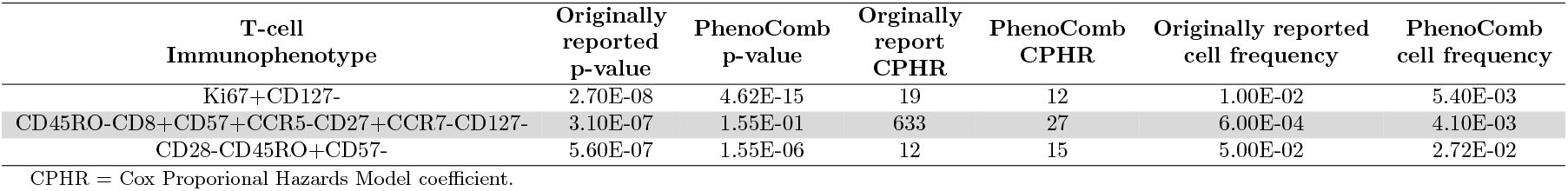
HIV phenotype evaluation comparison.

| T-cell<br>Immunophenotype | Originally<br>reported<br>p-value | PhenoComb<br>p-value | Originally<br>report<br>CPHR | PhenoComb<br>CPHR | Originally reported<br>cell frequency | PhenoComb<br>cell frequency |
| --- | --- | --- | --- | --- | --- | --- |
| Ki67+CD127- | 2.70E-08 | 4.62E-15 | 19 | 12 | 1.00E-02 | 5.40E-03 |
| CD45RO-CD8+CD57+CCR5-CD27+CCR7-CD127- | 3.10E-07 | 1.55E-01 | 633 | 27 | 6.00E-04 | 4.10E-03 |
| CD28-CD45RO+CD57- | 5.60E-07 | 1.55E-06 | 12 | 15 | 5.00E-02 | 2.72E-02 |

CPHR = Cox Proportional Hazards Model coefficient.

We also benchmarked PhenoComb’s performance in assessing this dataset. For the 12 markers assessed, a total of 220,170 phenotypes were identified using a minimum cell count threshold of 100 in 0.5 proportion of samples out of a theoretical maximum of 531,440 phenotypes (Fig. 2a). Computation time was 2.35 minutes (Fig. 2b).

### 3.3 Idenatification of Phenotypes Associated with COVID Infection

We next utilized PhenoComb on a publicly available CyTOF dataset. PhenoComb was utilized to discretize markers, assess combinations and compare frequencies between COVID positive and negative samples by Man-Whitney U tests. Given the computational time needed to assess 40 markers with multiple discrete states, we divided the dataset into four lineages and assessed a subset of markers in each independently (Supplemental Table 1). These lineages were: CD45+CD3+ (T-cells, 26 additional markers assessed), CD45+CD19+ (B-cells, 22 markers), CD45+CD56+ (NK cells, 16 markers), CD45+CD14+ (monocytes, 16 markers), and CD45+CD3-CD19-CD56-CD14- (other immune cells, 18 markers). The following number of phenotypes with an unadjusted p-value of less or equal than 0.0005 were identified for the corresponding parent lineage: CD45+CD3+, 437,751,153 phenotypes; CD45+CD19+, 46,171,572 phenotypes; CD45+CD56+, 87,828 phenotypes; CD45+CD14+, 588,011 phenotypes; and CD45+CD3-CD19-CD56-CD14-, 12625878 phenotypes. After filtering for independent phenotypes, the following number of phenotypes were identified for each parent lineage: CD45+CD3+, 8 phenotypes; CD45+CD19+, 13 phenotypes; CD45+CD56+, 21 phenotypes; CD45+CD14+, 15 phenotypes; and CD45+CD3-CD19-CD56-CD14-, 5 phenotypes.

Phenotypes with a confidence score of 1 are reported in Table 2 and graphs of the data are shown in Figure 3. Both the CD45+CD3-CD19-CD56-CD14- and the CD3+CD19+ population had elevated frequencies of cells expressing the proliferation marker Ki67 and high levels of the activation marker/ectoenzyme CD38 [9] in COVID positive samples (Figures 3a–3c). In T-cells, decreased CD4 cells expressing CD45RA and negative for expression of CCR7 and CD25 were observed in COVID positive samples relative to COVID negative(Figure 3d). In NK cells, elevated frequencies of cells expressing the proliferation marker Ki67 were seen in COVID positive samples (Figure 3e), and elevated frequencies of cells negative for Ki67 were seen in COVID negative samples (Figure 3f). Monocytes also had elevated frequencies of Ki67 expressing cells in COVID positive samples (Figure 3g). Increases in frequencies of the monocyte parent lineage expressing the dendritic cell marker CD141 (Figure 3h) and, separately, CD11b (Figure 3i) were also observed in COVID positive samples.

**Figure 3:**
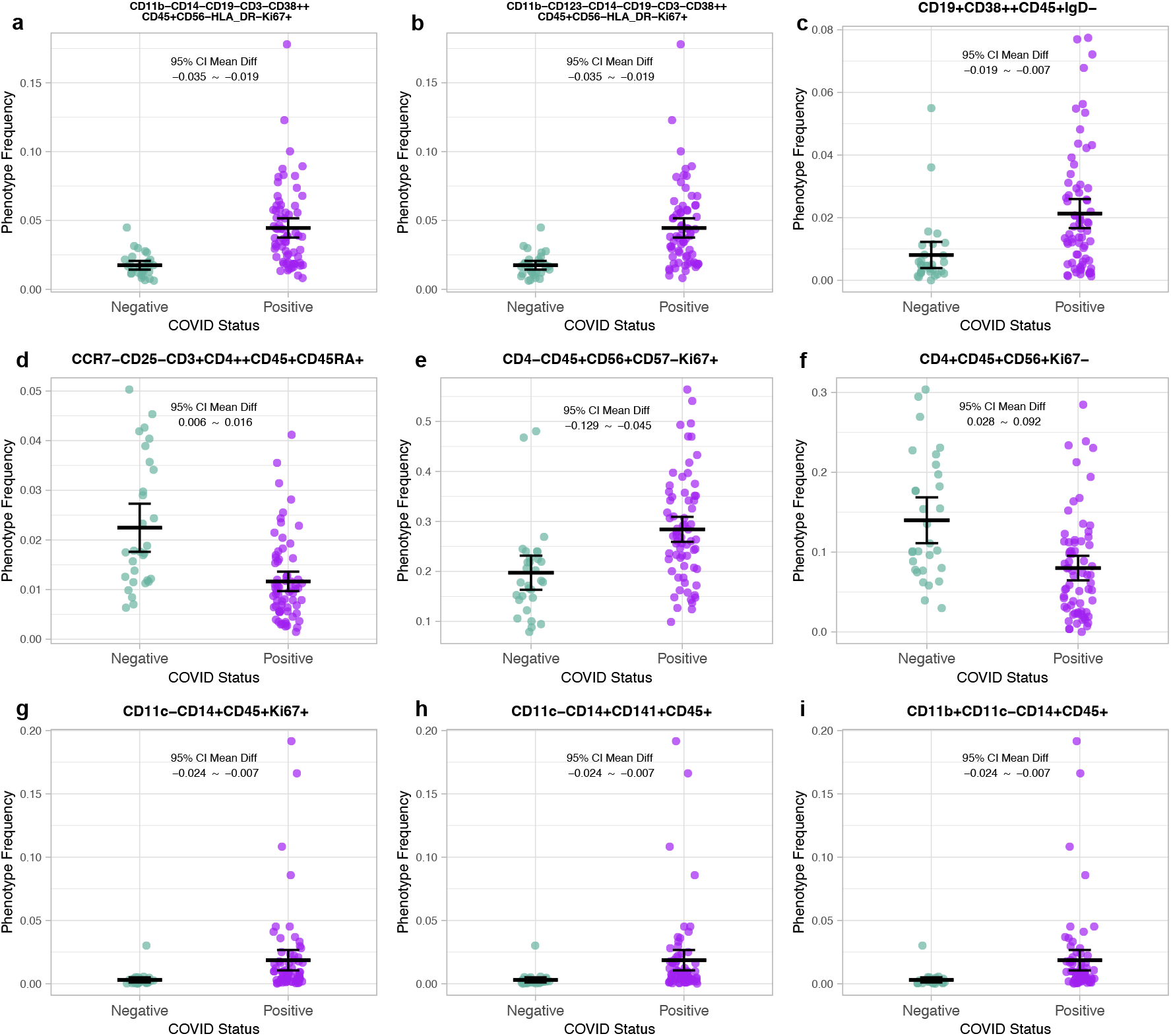
Identified Phenotypes. Phenotypes differing between COVID positive and COVID negative samples identified as independent with a confidence score of 1 are shown for the following parent lineages: (a,b) CD45+CD14-CD19-CD3-CD56-, (c) CD45+CD19+, (d) CD45+CD3+, (e,f) CD45+CD56+, (g,h,i) CD45+CD14+. Each panel shows the corresponding difference of means and the 95% confidence intervals.

**Table 2:** COVIDome identified phenotypes.

| Lineage | Phenotype | Standardized<br>Effect Size | Fold Change<br>(log<sub>2</sub>) |
| --- | --- | --- | --- |
| CD45+CD3-CD19-CD56-CD14- | CD11b-CD14-CD19-CD3-**CD38**++**CD45**+CD56-HLA_DR-**Ki67**+ | 0.71 | 1.35 |
| CD45+CD3-CD19-CD56-CD14- | CD11b-CD123-CD14-CD19-CD3-**CD38**++**CD45**+CD56-HLA_DR-**Ki67**+ | 0.71 | 1.35 |
| CD45+CD19+ | **CD19**+**CD38**++**CD45**+IgD- | 0.52 | 1.40 |
| CD45+CD3+ | CCR7-CD25-**CD3**+**CD4**++**CD45**+**CD45RA**+ | -0.57 | -0.95 |
| CD45+CD56+ | CD4-**CD45**+**CD56**+CD57-**Ki67**+ | 0.54 | 0.53 |
| CD45+CD56+ | **CD4**+**CD45**+**CD56**+**Ki67**- | -0.47 | -0.81 |
| CD45+CD14+ | CD11c-**CD14**+**CD45**+**Ki67**+ | 0.61 | 2.61 |
| CD45+CD14+ | CD11c-**CD14**+**CD141**+**CD45**+ | 0.61 | 2.61 |
| CD45+CD14+ | **CD11b**+CD11c-**CD14**+**CD45**+ | 0.61 | 2.61 |

For each of the lineages assessed, we also determined the total number of phenotypes present and the total runtime needed to compute. These data are plotted in Figures 2a and 2b. The following number of phenotypes were identified for the corresponding lineage (minimum cell count 100 in 50% of samples): CD45+CD56+, 485,832 phenotypes; CD45+CD14+, 1,537,542 phenotypes; CD45+CD3-CD19-CD56-CD14-, 28,219,848 phenotypes; CD45÷CD19÷, 184,492,651 phenotypes; and CD45+CD3+, 8,263,420,408 phenotypes. The lineages took the following times to compute:

CD45+CD56+, 17 minutes; CD45+CD14+, 18.37 minutes; CD45+CD3-CD19-CD56-CD14-, 18.53 minutes; CD45+CD19+, 3.26 hours; and CD45+CD3+, 49.97 hours.

## 4 Discussion

Herein we demonstrated that PhenoComb is an efficient and effective discovery tool for agnostically interrogating high-dimension cytometry data. The package pipeline is carried out over four steps: 1) discretizing marker expression based on user defined thresholds, 2) assessing the frequencies of all maker combinations present in the dataset for each sample, 3) performing a statistical comparison of the phenotype frequencies and 4) network/clustering analysis to identify representative independent phenotypes from those identified as significantly different. While we presented a pipeline analysis of data using PhenoComb, dataframes/csv files are generated at intermediate points that can be used for further interrogation inside and outside of PhenoComb. For example, the frequency of all identified phenotypes can be accessed to further interrogate specific phenotypes of interest, even if they are not identified as different in comparisons or an independent phenotype in clustering.

The presented analyses utilized two-state (i.e. +/-) or three state (i.e. negative, low, high) discretization of markers in datasets, but PhenoComb is theoretically compatible with a large number of discretely defined states. Defining the thresholds and the number of states is determined by the user, as it is in manual analysis. The number of states should be guided by visual analysis and background knowledge for markers. For example, CD4 is expressed by monocytes (and other innate cell lineages) in addition to T-cells, with monocytes having a “low” expression state and T-cells having a “high” expression state. Hence, thresholds for CD4 should define three states if the staining index is adequate to discriminate. Caution should be taken not to define more states than are supported by the biology of the marker and the quality of staining. While measurements for markers in cytometry are continuous, much of the variation is technical in nature rather than biological.

While we generated synthetic datasets to assess the computational time necessary, real datasets take significantly less time to calculate than the theoretical. As mentioned before and shown in our analysis of the HIV and COVIDome datasets, this is due to many of the theoretical marker combinations being absent in real data. This is the consequence of a limited number of cells being interrogated and/or biological limitations to co-expression/exclusion of markers. With a large enough cell count per sample, an interesting question that PhenoComb is capable of answering is how many of the theoretical marker combinations are present. In other words, PhenoComb could be applied to identify if and what phenotypes are absent. Likewise, PhenoComb could be used to interrogate highly correlated markers, which then could be used to inform more efficient future analyses.

Using PhenoComb to assess the HIV and COVIDome datasets, we demonstrated it was able to process large, real-world datasets in a relatively short amount of time. For example, in the COVIDome dataset analysis of the CD45+CD14+ lineage, PhenoComb was able to assess over 1.5 million phenotypes derived from 16 markers across 99 samples in under 19 minutes. However, given the super-exponential increase in computation time, increases in the maximum length of phenotypes can result in analyses requiring extended amounts of time. Illustrating this, the CD45+CD3+ lineage, also from the COVIDome dataset, took 50 hours to assess 26 markers. While for specialized and/or occasional use this may be an acceptable amount of time, often it is impractical. This can be mitigated through tuning functional arguments in PhenoComb, including filtering out low frequency populations or limiting the maximum length of phenotypes to less than the total number of markers interrogated.

In interrogating the HIV dataset, PhenoComb identified two of the three phenotypes associated with time to seroconversion highlighted in the original publication. However, there was variation between the p-values, Cox Proportional Hazard coefficients and cell frequencies between PhenoComb and the original publication. This likely reflects the differences in statistical tests utilized and differences caused by the subjective nature of gating markers. This likely also explains why PhenoComb did not identify one of the three original publication phenotypes as significant.

In the COVIDome dataset, PhenoComb identified Ki67+ phenotypes in the NK cell, monocyte and undefined other immune cell (i.e. CD45+CD3-CD19-CD56-CD19-) lineages elevated in peripheral blood samples from COVID positive individuals. Increases in B-cells and undefined other immune cells expressing high levels of CD38 were also observed. These phenotypes highlight proliferation and activation of immune cells as elevated in COVID infection. These phenotypes are associated with an active immune response, as would be expected in an infected individual. In T-cells, a CD4++CD45RA+CCR7-CD25-phenotype was decreased in COVID positive individuals. A CD45RA+CCR7-T-cell phenotype is canonically associated with an effector or “effector memory re-expressing RA (TEMRA)” phenotype [10, 11]. A decrease in effector T-cells in COVID infected individuals is counter-intuitive; however, the lack of CD25 expression suggests that these cells are not activated [12] and hence not truely effector cells.

It is important to note that for all statistical tests, p-values are not corrected for the number of comparisons. This is because each phenotype assessed is not an independent observation of the data. Instead, a large number of phenotyes are dependent on each other, making it difficult to estimate multiple comparison adjusted p-values. Inflation of type I errors is partially mitigated by the network/clustering approach that uses metrics other than p-values to rank their importance. Regardless, identified phenotypes should be treated as “discoveries”, requiring further validation.

PhenoComb is a powerful tool to add to the cytometrist’s analysis toolset. By taking full advantage of the single cell, high-dimension data generated in cytometry assays, PhenoComb empowers exploratory data analysis and is able to highlight phenotypes for further characterization. It’s im-plementation is also flexible, making it adaptable to work with various single-cell data types.

## Acknowledgements

We extended our appreciation to Eric Clambey, director of the Flow Cytometry Shared Resource at the University of Colorado Cancer Center, for his assistance with the CyTOF dataset. We also extend our appreciation to the original authors of the HIV and COVIDome datasets and to Flow Repository for their efforts in supporting open science.

---

**Supplemental Table 1.** Lineages from COVIDome dataset.

|  | CD45+CD3+ | CD45+CD19+ | CD45+CD56+ | CD45+CD14+ | CD45+CD3-CD19-CD56-CD14- |
| --- | --- | --- | --- | --- | --- |
| CD45 |  |  |  |  |  |
| CD57 | X | X | X | X | X |
| CD11c |  | X |  | X | X |
| CD16 |  | X | X | X | X |
| CD196 CCR6 | X | X | X | X | X |
| CD19 |  |  |  |  |  |
| CD123 | X | X | X | X | X |
| CCR5 | X | X | X | X | X |
| IgD |  | X |  |  | X |
| CD1c |  | X |  | X | X |
| CD38 | X | X | X | X | X |
| CD127 | X |  |  |  |  |
| CD86 |  | X |  | X | X |
| ICOS | X | X |  |  |  |
| CD141 |  |  |  | X | X |
| TGIT | X |  | X |  |  |
| Tim3 | X |  |  |  |  |
| CD27 | X |  |  |  |  |
| CXCR3 | X |  |  |  |  |
| CD45RA | X | X |  |  |  |
| PD-1 | X | X |  |  |  |
| PDL1 | X | X | X | X | X |
| CD14 |  |  |  |  |  |
| Tbet | X | X | X |  |  |
| Ki67 | X | X | X | X | X |
| CD33 |  |  |  | X | X |
| CD95 | X | X |  |  |  |
| Foxp3 | X | X |  |  |  |
| Eomes | X |  | X |  |  |
| CCR7 | X |  |  |  |  |
| CD8a | X |  | X |  |  |
| CD25 | X | X |  |  |  |
| CD3 |  |  | X |  |  |
| CXCR5 | X | X | X | X | X |
| IgM |  | X |  |  | X |
| HLA-DR | X | X | X | X | X |
| CD4 | X |  | X |  |  |
| CCR4 | X |  |  |  |  |
| CD56 |  |  |  |  |  |
| CD11b |  |  |  | X | X |
| **Total markers** | **26** | **22** | **16** | **16** | **18** |

\*X = included in analysis

**Supplemental Figure 1:**
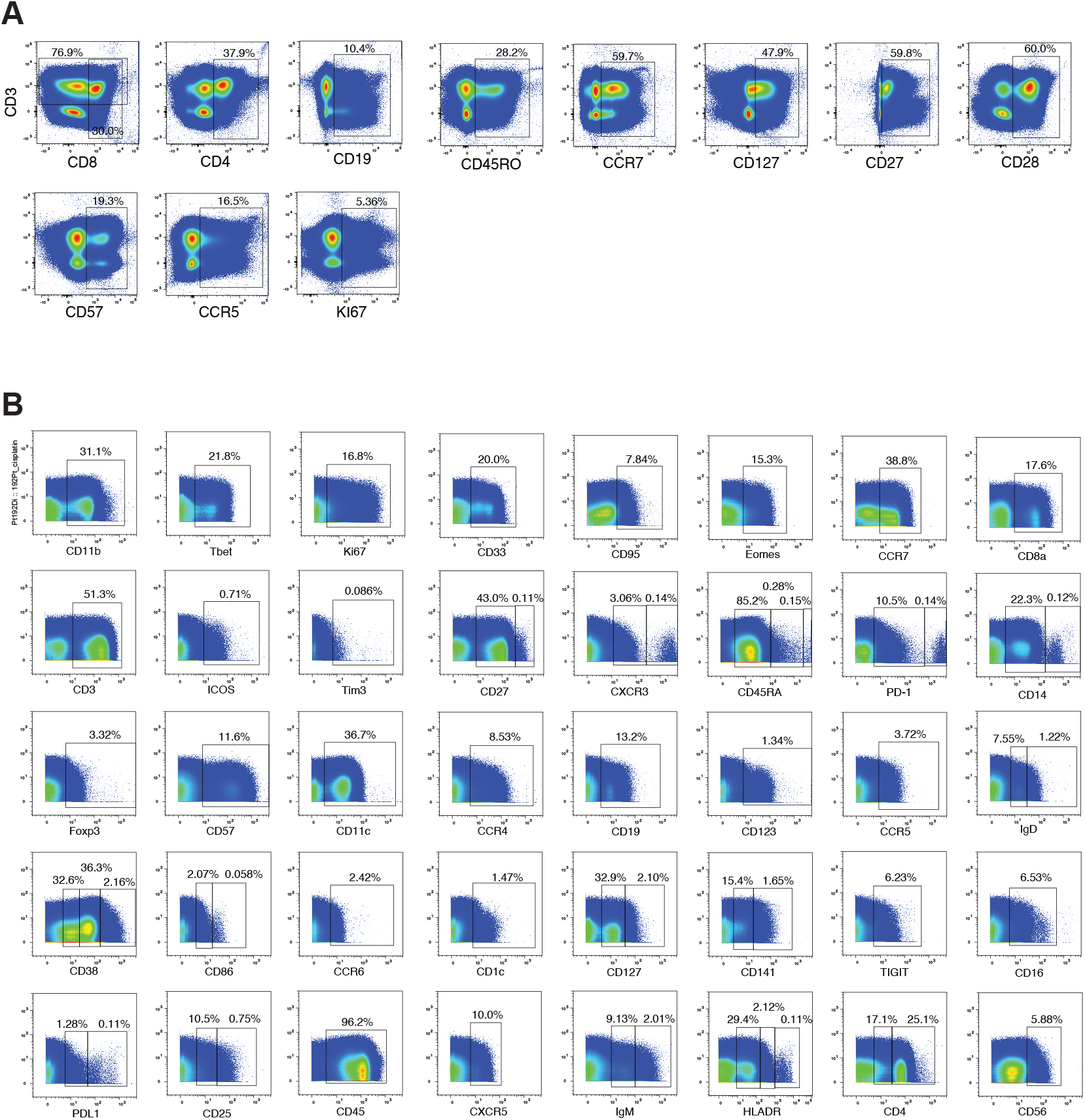
Gating Strategies. Gating strategies for the **(A)** HIV dataset and for **(B)** COVIDome dataset. The rectangle’s limits on the x axis are used as thresholds for obtaining discrete marker states.

## References

[1] Pratip K. Chattopadhyay, Carl-Magnus Hogerkorp, and Mario Roederer. A chromatic explosion: the development and future of multiparameter flow cytometry. Immunology, 125(4):441–449, December 2008.

[2] Etienne Becht, Leland McInnes, John Healy, Charles-Antoine Dutertre, Immanuel W H Kwok, Lai Guan Ng, Florent Ginhoux, and Evan W Newell. Dimensionality reduction for visualizing single-cell data using UMAP. Nature Biotechnology, 37(1):38–44, January 2019.

[3] Jacob H. Levine, Erin F. Simonds, Sean C. Bendall, Kara L. Davis, El-ad D. Amir, Michelle D. Tadmor, Oren Litvin, Harris G. Fienberg, Astraea Jager, Eli R. Zunder, Rachel Finck, Amanda L. Gedman, Ina Radtke, James R. Downing, Dana Pe’er, and Garry P. Nolan. Data-Driven Phenotypic Dissection of AML Reveals Progenitor-like Cells that Correlate with Prognosis. Cell, 162(1):184–197, July 2015.

[4] David M. Woods, Andressa S. Laino, Aidan F. Winters, Jason M. Alexandre, Daniel Freeman, Vinay Rao, Santi S. Adavani, Jeffrey S. Weber, and Pratip K. Chattopadhyay. Nivolumab and ipilimumab are associated with distinct immune landscape changes and response-associated immunophenotypes. JCI Insight, May 2020.

[5] Nima Aghaeepour, Pratip K. Chattopadhyay, Anuradha Ganesan, Kieran O’Neill, Habil Zare, Adrin Jalali, Holger H. Hoos, Mario Roederer, and Ryan R. Brinkman. Early immunologic correlates of HIV protection can be identified from computational analysis of complex multivariate T-cell flow cytometry assays*. Bioinformatics, 28(7):1009–1016, April 2012.

[6] Vincent D Blondel, Jean-Loup Guillaume, Renaud Lambiotte, and Etienne Lefebvre. Fast unfolding of communities in large networks. Journal of Statistical Mechanics: Theory and Experiment, 2008(10):P10008, October 2008.

[7] Amy C Weintrob, Ann M Fieberg, Brian K Agan, Anuradha Ganesan, Nancy F Crum-Cianflone, Vincent C Marconi, Mollie Roediger, Susan L Fraser, Scott A Wegner, and Glenn W Wortmann. Increasing Age at HIV Seroconversion From 18 to 40 Years Is Associated With Favorable Virologic and Immunologic Responses to HAART. JAIDS Journal of Acquired Immune Deficiency Syndromes, 49(1):40–47, September 2008.

[8] Kelly Daniel Sullivan, Matthew Dominic Galbraith, Kohl Thomas Kinning, Kyle William Bartsch, Nik Caldwell Levinsky, Paula Araya, Keith Patrick Smith, Ross Erich Granrath, Jessica Rose Shaw, Ryan Michael Baxter, Kimberly Rae Jordan, Seth Aaron Russell, Monika Ewa Dzieciatkowska, Julie Ann Reisz, Fabia Gamboni, Francesca Isabelle Cendali, Tusharkanti Ghosh, Andrew Albert Monte, Tellen Demeke Bennett, Michael George Miller, Elena Wen-Yuan Hsieh, Angelo D’Alessandro, Kirk Charles Hansen, and Joaquin Maximiliano Espinosa. The COVIDome Explorer researcher portal. Cell Reports, 36(7):109527, August 2021.

[9] Valeria Quarona, Gianluca Zaccarello, Antonella Chillemi, Enrico Brunetti, Vijay Kumar Singh, Enza Ferrero, Ada Funaro, Alberto L. Horenstein, and Fabio Malavasi. CD38 and CD157: A long journey from activation markers to multifunctional molecules: CD38 and CD157. Cytometry Part B: Clinical Cytometry, 84B(4):207–217, July 2013.

[10] Jeffrey D. Ahlers and Igor M. Belyakov. Memories that last forever: strategies for optimizing vaccine T-cell memory. Blood, 115(9):1678–1689, March 2010.

[11] Nicholas P. Restifo and Luca Gattinoni. Lineage relationship of effector and memory T cells. Current Opinion in Immunology, 25(5):556–563, October 2013.

[12] A. Caruso, S. Licenziati, M. Corulli, A. D. Canaris, M. A. De Francesco, S. Fiorentini, L. Peroni, F. Fallacara, F. Dima, A. Balsari, and A. Turano. Flow cytometric analysis of activation markers on stimulated T cells and their correlation with cell proliferation. Cytometry, 27(1):71–76, January 1997.

